# Retrieval-related eye movements are predictive of memory precision

**DOI:** 10.64898/2025.12.17.694913

**Authors:** Mingzhu Hou, Luke R. Pezanko, Sabina Srokova, Paul F. Hill, Arne D. Ekstrom, Michael D. Rugg

## Abstract

Eye movements are predictive of successful episodic memory encoding and retrieval, but it is unclear whether they reflect the precision of retrieved memory content. Here, we examined relationships between eye fixations, memory precision and fMRI BOLD activity. At study, participants were presented with object images, each placed at a random location on an invisible circle. At test, both studied and new images were presented. Participants were instructed to make a covert recognition memory judgment to each image, recall and then signal its studied location, guessing if necessary. Simultaneous fMRI and eye-tracking data were acquired during the test phase. For correctly recognized images, the trial-wise angular distance between the studied and reported locations was fitted to a two-component mixture model. Based on the model-derived parameters, correctly recognized images were categorized as those associated with successful location retrieval (location hits) or guesses. Location hits were further divided into high- and low precision trials. At retrieval, fixations predicted both successful retrieval and the precision of the retrieved location memory. Additionally, across-trial fixation patterns were more similar for location hits than for guesses, and for high- than for low-precision trials. The association between retrieval success and eye fixation precision was evident across almost the entire recall phase, whereas memory precision effects emerged around 1-1.5s after image onset. Fixation precision and fixation pattern similarity each predicted trial-wise memory precision independently of hippocampal BOLD activity, which also predicted precision. These findings suggest that eye fixations during retrieval track the fidelity of mnemonic content.

## Introduction

Episodic memory is mainly assessed through overt memory judgments, but analysis of incidental eye movements can provide additional insight into how episodic memories are encoded and retrieved. Notably, fixation patterns during both the study and test phases of episodic memory tasks have been reported to reflect mnemonic content (for reviews, see Ryan & Shen, 2020; Wynn et al., 2019). Here, we focus on the relationship between fixation patterns and memory performance at retrieval.

Prior findings provide evidence for an association between fixation patterns and memory retrieval (e.g., Damiano & Walther, 2019; Hannula et al., 2009; Johansson & Johansson, 2014; for reviews, see Hannula et al., 2010; Ryan & Shen, 2020). For example, in a study examining object-location memory, Johnasson and Johnasson (2014) reported that in a free-viewing condition, fixations were more likely to be directed toward the quadrant of the display where object images had been studied. By contrast, constraining eye movements to a different location impaired memory for the object. The findings from this and other studies cited above indicate that fixation patterns during retrieval predict successful memory retrieval.

Other recent studies have examined associations between anticipatory gaze and retrieved mnemonic content. For instance, Schmidig et al. (2025) reported that when participants re-watched animated movie clips containing a surprising event, they made anticipatory fixations toward the location of the event prior to its occurrence. Furthermore, anticipatory fixations that were closer to the event’s location on the display were associated with an increased likelihood of later identifying the quadrant of the screen in which the event occurred. Similarly, Buchel et al. (2024) had participants repeatedly learn a list of image exemplars belonging to one of five visual categories. Across repetitions, participants showed improved memory performance for the sector containing each exemplar and demonstrated anticipatory fixations that shifted closer to the exemplar’s location.

The findings from the aforementioned studies suggest that fixation patterns at retrieval vary according to the content of the retrieved memory, but they do not speak to the question of whether fixations are predictive of memory *precision*. Addressing this question will shed light on the fidelity with which eye movements track retrieved memory content and their potential as an incidental marker of successful retrieval (cf., Schmidig et al., 2025). Recent studies examining memory precision have employed a positional response accuracy task (Harlow & Donaldson, 2013), in which memory performance is evaluated with a continuous metric that reflects the degree to which a feature of the study event can be accurately reproduced. Findings from behavioral studies employing positional response accuracy tasks suggest that retrieval success is dissociable from memory precision (e.g. Gellerson et al., 2024; Harlow & Yonelinas, 2016; see also Kolarik et al., 2018 and Nilakantan et al., 2018 for evidence from patients with hippocampal damage).

Several studies have employed functional magnetic resonance imaging (fMRI) to examine the neural correlates of retrieval success and precision, consistently implicating the hippocampus and left angular gyrus (AG) (Richter et al., 2016; Cooper et al., 2017; Korkki et al., 2023; Hou et al., 2025a, b). In a recent study (Hou et al., 2025b), we examined the fMRI correlates of memory precision in a combined dataset that included the same sample we report on below as well as a separate sample that was originally described by Hou et al. (2025a). We reported that BOLD activity in the hippocampus and the left AG tracked trial-wise estimates of memory precision, whereas no effects of retrieval success could be identified in either region.

The present study had two primary aims. The first aim was to examine whether eye fixations predict the precision with which location information can be retrieved. Given prior evidence demonstrating robust associations between eye movements and neural activity in the hippocampus (for reviews, see Kragel & Voss, 2022; Ryan et al., 2020), our second aim was to examine associations between memory precision, precision-related eye fixations, and hippocampal fMRI BOLD signals.

## Materials & Methods

The behavioral and fMRI data pertaining to the present study sample were first reported as part of the dataset described by Hou et al. (2025b). Here, we report the relationships between these data and eye tracking metrics. These data have not been reported previously.

### Participants

Participants were 24 young adults (mean age = 23 yrs, age range = 18-33 yrs, 11 female). The sample size was consistent with that of prior fMRI studies examining the neural correlates of memory precision in young adults (Hou et al., 2025a, n = 23; Korkki et al., 2023, n = 23; Richter et al., 2016, n = 21) and with prior studies examining the relationship between eye fixations and memory performance (e.g., Buchel et al., 2025, n = 27; Johansson & Johansson, 2014, n = 24). All participants were cognitively healthy, right-handed, had normal or corrected-to-normal vision, no history of neurological or psychiatric illness, and were not taking prescription medications that affected the central nervous system.

Informed consent was obtained in accordance with the University of Texas at Dallas Institutional Review Board guidelines. Participants were compensated at the rate of $30 an hour. Eye movement data from two participants were excluded due to poor calibration, defined as a validation error exceeding 1 deg for any fixation as based on Eyelink’s default threshold. Accordingly, we report data from the remaining 22 participants.

### Experimental items

The experimental items were 136 images of everyday objects (see Figure 1). 102 of these images were used as study items and the remaining 34 images served as new (unstudied) test items.

**Figure 1.**
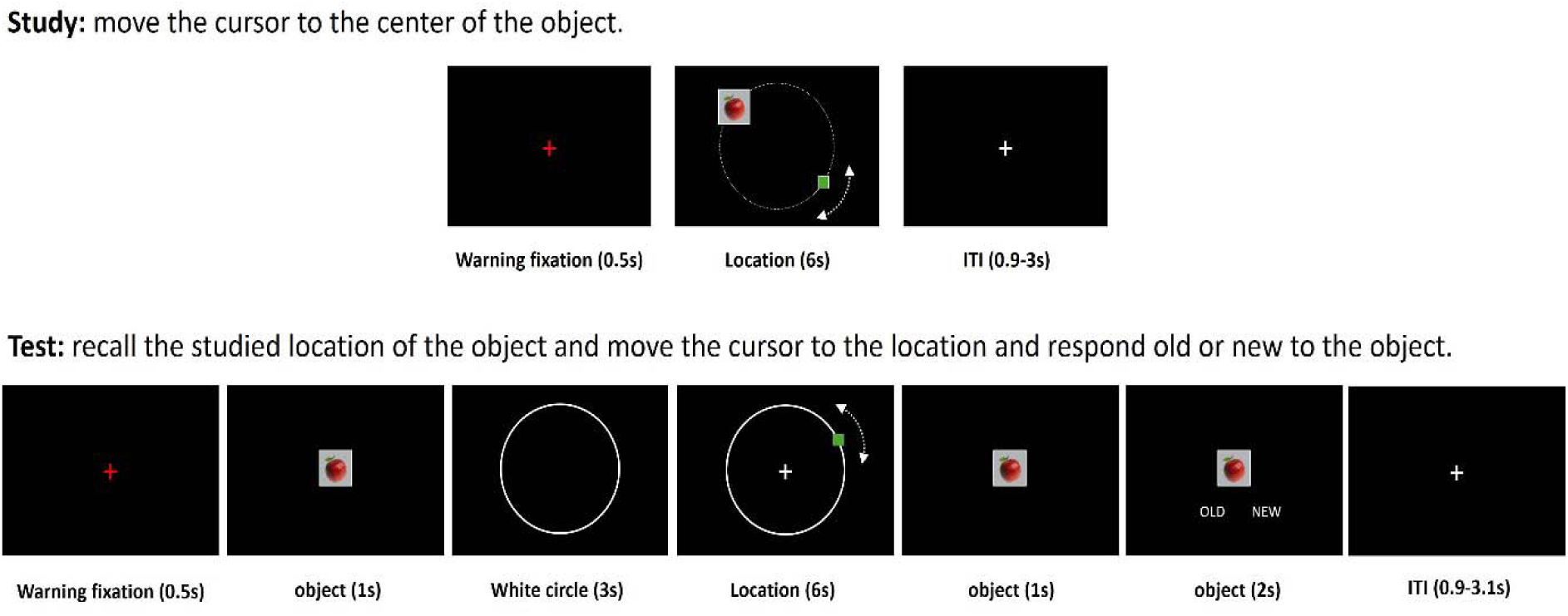
Schematics of the study and test phases.

### Procedure

The schematics for the experimental tasks are shown in Figure 1. There was a single study block followed by four consecutive test blocks. Both the study and test phases were undertaken in the MRI scanner.

At study, participants viewed 102 object images (visual angle = 1.4 x 1.4 deg, image size = 123 x 123 pixels), each presented for 6 s at a random location on a virtual circle (radius = 4.9 deg visual angle, equivalent to 422 pixels). Along with the image, a small green cursor was presented on the circle at a distance of at least 60 deg from the image. The instruction was to use a button box to move the cursor around the circle to the center of the image, and to remember both the image and its location. A 30-s break occurred in the middle of the study block, during which a ‘rest’ cue was presented at the center of the screen. Participants were instructed to relax until the disappearance of the cue.

The test images comprised all the studied images intermixed with 34 unstudied images. They were presented pseudo-randomly so that no more than 3 studied or new images occurred in succession. Each test trial began with a presentation of the test image for 1 s, after which a white circle of the same size as the virtual circle used during the study phase was presented for 3 s. Participants were instructed to covertly recall the studied location of the image. They were then to move a cursor that was randomly located on the circle (with the constraint that it was no less than 60 deg from the image’s studied location) to the recalled location, again by using a button box, guessing if necessary. After moving the cursor, participants responded whether the presented object to have been studied or unstudied.

Participants viewed the experimental stimuli via a mirror mounted on the scanner head coil that reflected a back-projected image (viewing distance 102 cm). They pressed the buttons under their right index and middle fingers to move the cursor along the circle during the study and test phases, and to respond ‘old’ or ‘new’ to the test image. The mappings between the button presses and the cursor movement direction, as well as the old/new judgments, were counterbalanced across participants.

Eye movement data were acquired during the study and test phases with an MRI-compatible eye-tracker (Eyelink 1000 Plus; SR Research). Monocular data were collected from the right eye at a sampling rate of 1000 Hz. Since drift in eye-tracking accuracy is minimal when calibration is conducted just prior to a scanner run (Peitek et al., 2018), calibration and manual validation were performed using Eyelink’s 9-point calibration procedure immediately before each run. This procedure was repeated until the validation accuracy at each gaze point was acceptable according to Eyelink’s default threshold (1 deg). We used Eyelink’s default event parser to classify eye movements into saccades, blinks and fixations. Saccades were defined by a velocity threshold of 30 deg/s and an acceleration threshold of 8000 deg/s^2^. Samples with missing pupil size data were classified as blinks. The remaining data were categorized as fixations.

### Behavioral analyses

#### Item memory performance

Item memory performance (Pr) was calculated as the difference between the proportion of item hits (studied objects that were correctly recognized) and the proportion of false alarms to new items.

#### Separation of location hits and guess trials

For each item hit trial, distance error was calculated as the angular difference between the reported location and the studied location. Distance errors for item hits were pooled across all participants and fitted to a mixture model using the MLE() function implemented in MemToolbox (Suchow et al., 2013). This procedure estimates a rectangular distribution of the proportion of guess responses (g), and a von Mises distribution of item hit trials associated with varying levels of memory precision [quantified by the concentration parameter (Kappa)].

Item hit trials were divided into successful (location hit) and guess trials depending on whether or not the associated distance errors had a < 5% chance of fitting the aforementioned across-participant von Mises distribution (estimated with the “vonmisecdf” function from www.paulbays.com/toolbox/). According to this criterion, the cutoff for distinguishing location hits from guess trials was +/- 36 deg. Therefore, location hits were defined as item hit trials with an absolute distance error < 36 deg, and guess trials were defined as those with an absolute distance error of between 36 and 180 deg.

To facilitate the comparison between location hit trials with varying levels of precision, the trials were subdivided into high-precision and low-precision trials based on their associated absolute distance errors. As in our prior studies (Hou et al., 2025a, b), high-precision trials were defined as location hits with an absolute distance error < 16 deg. Low-precision trials were defined as location hits with an absolute distance error ranging from 20-35 deg. The small gap between these ranges was to reduce the possibility of misclassifying trials that may have been more appropriately categorized as the alternate trial type.

### Analyses of eye fixation data

#### Preprocessing of fixation data

Eye movement data were preprocessed using the edfmex.m function in MATLAB 2023a (The MathWorks, Inc. 2023). Fixations with a duration shorter than 50 ms were excluded from analyses. To differentiate image-related fixations from location-related fixations, a square was drawn at the center of the screen with sides twice the length of the test image. Location-related fixations (hereafter, fixations) were defined as those falling outside this central square.

#### Effects of retrieval success and memory precision on fixation metrics

In the following analyses, we derived trial-wise measures of the distance between fixation location and the location of the image at study (hereafter, fixation precision). Additionally, we computed the similarity of fixation distribution patterns separately for location hits and guess trials (i.e., fixation pattern similarity, see below). In these analyses, retrieval success effects were examined by contrasting these fixation metrics between location hits and guess trials. Memory precision effects were operationalized as differences in fixations metrics according to the precision of location hit trials.

#### The effect of retrieval success on fixation precision

For a given item hit trial, fixation precision was operationalized as the mean Euclidean distance (in pixels) of each fixation relative to the studied location during the 4s interval extending from the onset of the test image to the end of the covert recall phase. We elected to employ Euclidean distance rather than angular error when estimating fixation precision to allow for the fact that fixations might be directed to locations above or below, as well as along, the circle on which the image was located at study. To assess differences in fixation precision between location hits and guess trials, we conducted a pairwise t-test comparing the fixation precision metrics for the two classes of trial across participants.

#### The effect of memory precision on fixation precision

To examine whether eye fixation metrics were sensitive to memory precision, we employed linear mixed-effects (LME) analyses using the lmer() function in R (R Core Team, 2018). An LME model was constructed to examine the relationship across location hits between trial-wise fixation precision and trial-wise judgment precision (measured by absolute distance error). The model employed fixation precision as a fixed effect predictor of trial-wise absolute distance error. Participants were modeled as a random intercept [LME syntax: memory precision ∼ fixation precision + (1|participant)].

#### Time series analyses

To examine the time course of the fixations associated with location hit and guess trials, we conducted a time-series analysis, in which the 4-s recall phase was separated into 8 successive 500-ms time bins. Fixation precision in each time bin was calculated as the precision of each fixation falling within the bin, weighted by its duration within that bin. An LME model was implemented with trial type (location hit, guess), time bin and their interaction as the predictors of fixation precision. Participants were treated as a random intercept [LME syntax: fixation precision ∼ trial type + time bin + trial type x time bin + (1|participant)]. A significant trial type x time bin interaction was followed up with an LME analysis employing trial type (location hit vs. guess) to predict fixation precision at each time bin [LME syntax: fixation precision ∼ trial type + (1|participant)]. Because not all participants had fixations in all time bins, we implemented an LME approach rather than a participant-wise ANOVA because it is more applicable to the analysis of unbalanced data. We conducted an analogous analysis to compare the time courses of fixations associated with high vs. low precision location hits. In both sets of analyses, significant time bins were defined as those that survived false discovery rate (FDR) correction.

To examine whether any potential effect of trial type was associated with differences in pre-stimulus fixation patterns, we additionally analyzed fixation precision for the 500 msec time bin immediately prior to the onset of the test image (see Figure 1).

#### Pattern similarity analysis

To examine whether the spatial patterning of eye fixations varied with trial type, we conducted pattern similarity analyses on the fixation data. Because the studied locations were unique across the item hit trials, the locations were realigned to a common reference point (the top point of the virtual circle, see Figure 5). Coordinates of fixations within each trial were adjusted accordingly to preserve their relative distance to the studied location.

Pattern similarity analyses were performed using the eyesim package (Buchsbaum, 2025). For each participant, trial-wise fixation density maps were generated by applying a duration-weighted Gaussian smoothing kernel (sigma = 50 pixels) to the adjusted fixations. The density map for each trial was then vectorized by flattening the density values from all cells into a single column vector.

For each participant, we computed trial-wise pattern similarity metrics for location hits and guesses. For each location hit trial, pattern similarity was computed as the Fisher z-transformed correlation between its density map and the maps corresponding to all other location hit trials, minus the mean Fisher z-transformed correlation between its associated density map and the maps corresponding to all guess trials. An analogous similarity metric was computed for guess trials. We employed one-sample t-tests to test the similarity metrics associated with different trial types against the null hypothesis of zero similarity. Pairwise t-tests were conducted to compare the participant-wise similarity metrics between location hits and guesses, and between high and low-precision trials.

#### Control analysis

We examined whether eye fixations were merely a predictor of the location of the upcoming manual judgment (i.e., cursor placement) independent of the accuracy of the retrieved mnemonic content. This was achieved by repeating the aforementioned fixation precision and pattern similarity analyses relative to the location signaled by the participants (i.e., judged location) rather than the studied location. We compared the resulting ‘fixation precision’ metric between location hit and guess trials, and separately between high and low precision trials. Similarity metrics for the different trial types were contrasted against a null hypothesis of zero. They were also contrasted between location hit and guess trials and between high and low precision trials.

### MRI acquisition and preprocessing

As reported in more detail previously (Hou et al., 2025b), MRI data were acquired with a Siemens PRISMA 3T MR scanner equipped with a 32-channel head coil. Functional images were acquired with a T211-weighted echoplanar sequence [TR = 1560 ms, TE = 30 ms, flip angle = 70°, field-of-view (FOV) = 220 mm, multiband factor = 2, 48 slices, voxel size = 2.5 x 2.5 x 2.5 mm, 0.5 mm inter-slice gap, anterior-to-posterior phase encoding direction]. A T1-weighted image was acquired with a Magnetization-Prepared Rapid Acquisition Gradient Echo pulse sequence (TR = 2300 ms, TE = 2.41 ms, FOV 256 mm, voxel size = 1 x 1 x 1 mm, 160 slices, sagittal acquisition). A field map was acquired after the functional scans using a double-echo gradient echo sequence (TE 1/2 = 4.92 ms/7.38 ms, TR = 520 ms, flip angle = 60°, FOV = 220 mm, 48 slices, 2.5mm slice thickness).

Data preprocessing was performed with the SPM12 software package (Wellcome Department of Imaging Neuroscience, London, UK: www.fil.ion.ucl.ac.uk/spm) implemented in MATLAB 2023a. The functional images were field-map corrected, realigned, reoriented to the anterior commissure–posterior commissure line and spatially normalized to SPM’s MNI EPI template. The normalized images were resampled to 2.5 mm isotropic voxels (thereby interpolating across the inter-slice gap) and smoothed with a 6 mm Gaussian kernel. The functional data from the different test blocks were concatenated using the spm_fmri_concatenate.m function prior to entry into participant-wise GLMs. Anatomical images were normalized to SPM’s MNI T1 template.

### MRI analyses

For each participant, we estimated trial-wise BOLD activity from the test phase using a single-trial GLM that implemented the least squares all approach (Abdulrahman & Henson, 2016; Mumford et al., 2012). Each test event was modeled as a separate event of interest, with event-related neural activity modeled with a 4s duration boxcar onsetting simultaneously with the test image. Six motion regressors and four constants modeling the mean BOLD signal in the test blocks were included as covariates of no interest.

We employed the same hippocampal and left AG regions of interest (ROIs) as those employed in Hou et al. (2025b). The hippocampal ROI was defined as all voxels falling within the 5-mm radius of the peak precision effect identified by Hou et al. (2025b) (MNI coordinates: 35, -20, -18. For the left AG, we employed a combined ROI based on the peak precision effects reported by Korkki et al. (2023) and Richter et al. (2016) (MNI coordinates: -54, -66, 33 and -54, -54, 33). The AG ROI comprised all voxels falling within a 5-mm radius of each peak. Trial-wise parameter estimates were extracted by averaging over all the voxels in each ROI.

In light of our prior findings (see Introduction) that trial-wise BOLD activity in both the hippocampus and left AG was predictive of memory precision, we conducted LME analyses to examine the relationships between trial-wise BOLD activity in these ROIs and the precision of memory judgments. If an ROI demonstrated sensitivity to trial-wise variability in precision, it was employed in follow-up analyses to examine whether the relationship was moderated by the inclusion of trial-wise fixation metrics as an additional predictor.

### Relationships among memory precision, fixation metrics and neural activity

We employed LME analyses to examine whether fixation metrics predicted memory precision, and whether either metric could account for variance in memory precision estimates beyond that accounted for by trial-wise BOLD activity. Additional analyses were conducted to assess associations between the fixation metrics and fMRI BOLD activity.

Statistical analyses were conducted with R software (R Core Team, 2018). For the pairwise t-tests conducted at each time bin, the significance level was set at p < 0.05 after FDR correction. For all other analyses significance levels were set at p < 0.05 after family-wise correction for multiple comparisons using the Bonferroni procedure. Note that we did not include random slopes in our LME models because their inclusion led to singular fits in most models. For models that did not exhibit singular fits when random slopes were included, the results were essentially unchanged from the models where slopes were modelled as fixed effects.

## Results

### Memory performance

Across participants, the mean Pr was 0.65 (std = 0.18). The mixture model yielded a precision (Kappa) value of 10.76 and a guess rate (g) of 0.38.

### The effect of retrieval success on fixation precision

Figure 2 depicts fixation precision (as operationalized by Euclidian distance) for location hits and guesses. Location hit trials were associated with fixations that were significantly closer to the studied location [t(21) = 8.42, p < 0.001, Cohen’s d = 2.09].

**Figure 2.**
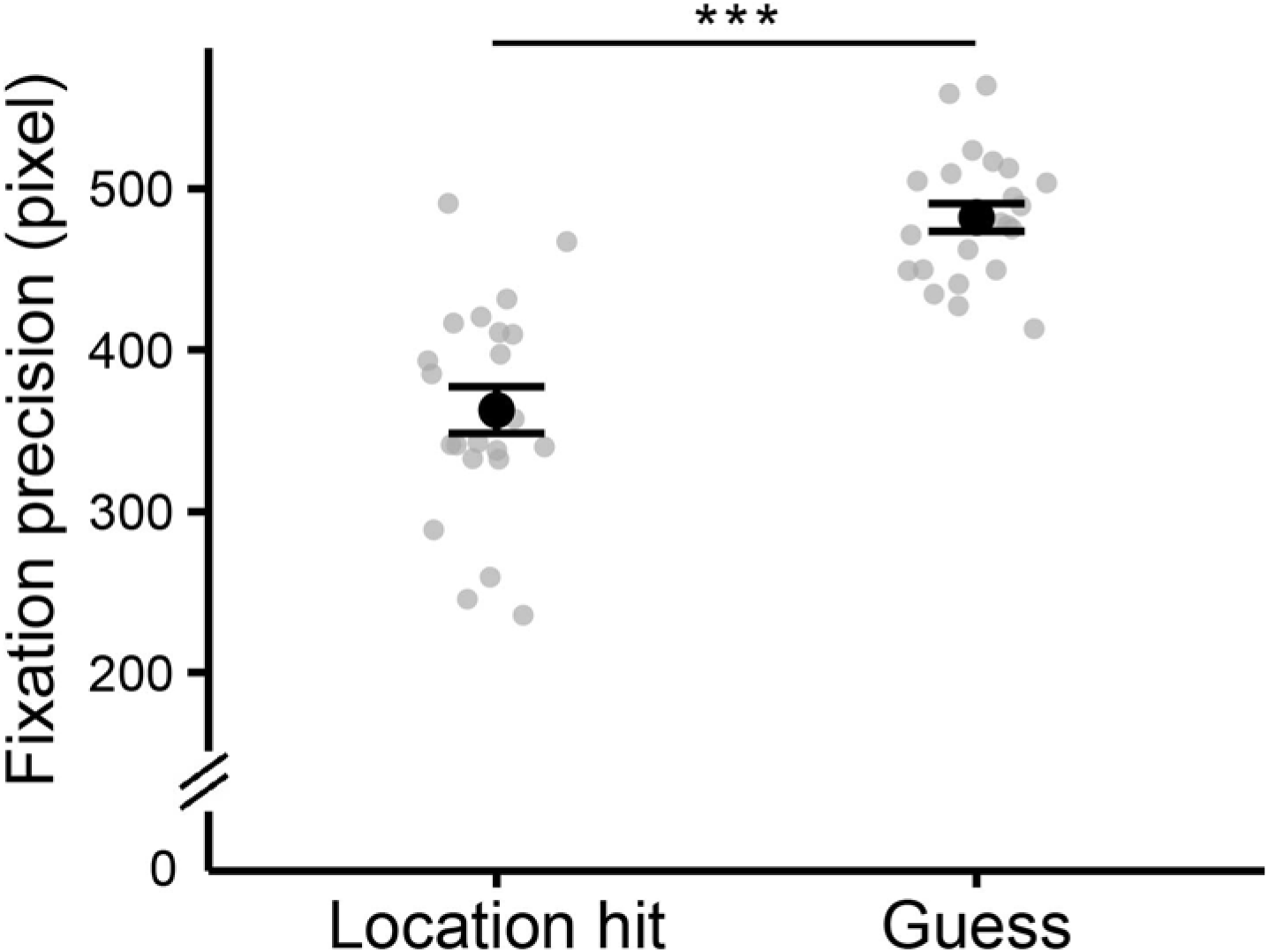
Fixation precision for location hit and guess trials. Error bars represent standard errors of the mean. The dots represent the participant-wise mean fixation to studied location distance. *** p < 0.001.

### Fixation precision was predictive of the precision of memory judgements

We examined the relationship between fixation precision and the precision of the corresponding location memory judgments, focusing on location hit trials. Table 1 shows the results from LME models that employed fixation precision as the predictor of the trial-wise precision of the corresponding memory judgment (indexed by abs. distance error). As is evident from the table, fixation precision was predictive of judgment precision; that is, fixations closer to the studied location were associated with more precise location judgments (Figure 3a).

**Figure 3.**
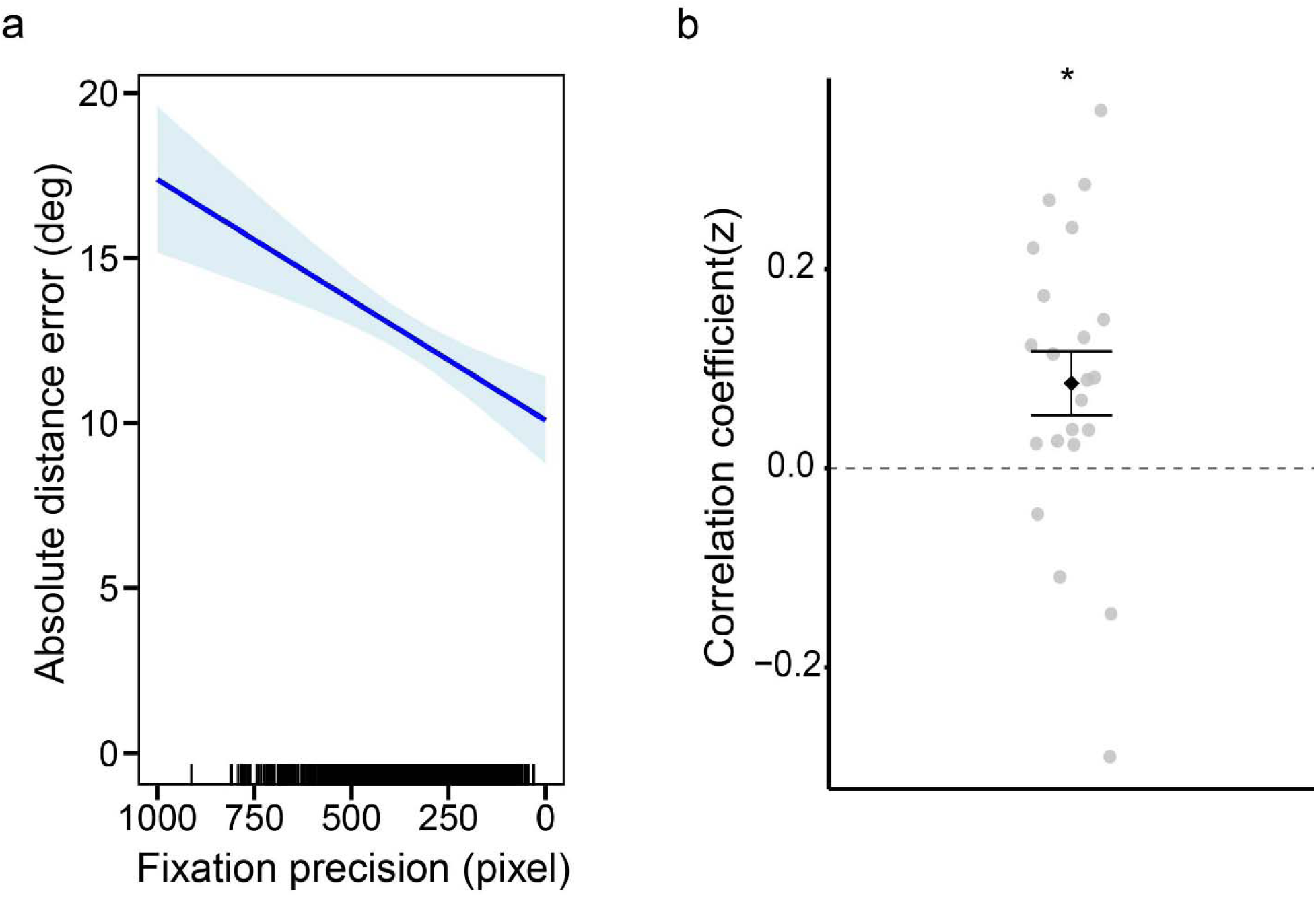
a: Plot depicting the relationship between trial-wise fixation precision and the model fitted values of judgment precision for location hits, as derived from the LME model employing fixation precision as the predictor of judgment precision. The shaded areas reflect the 95% confidence intervals. b: Participant-wise Pearson correlation coefficients (Fisher z-transformed) for the relationship across trials between fixation precision and judgment precision. The error bar represents the standard error of the mean. * p < 0.05.

**Table 1.** Results from a linear mixed model examining the relationship between fixation precision and trial-wise judgment precision for location hits. Standardized regression coefficients (β) are reported.

| Parameter | $\beta$ (SE) | df | t | p |
| --- | --- | --- | --- | --- |
| *Fixation-precision* |  |  |  |  |
| Fixation precision | 0.13 (0.03) | 584 | 4.35 | < 0.001 |

We also examined the pairwise correlation between the trial-wise angular error of the location judgments and fixation precision for each participant. The Fisher z transformed coefficients are shown in Figure 3b. Consistent with the findings from the LME model, a one-sample t-test revealed that the mean correlation coefficient was significantly above 0, t(21) = 2.67, p = 0.014, Cohen’s d = 0.57.

### Time series analyses

Table 2 shows the results from LME models examining the effects of trial type and time bin (500-ms bins, ranging from 0–4s) on fixation precision. As is evident from the table, the model comparing location hits and guess trials revealed significant main effects of trial type and time bin, as well as an interaction between the two factors. Follow-up analyses revealed that location hits were associated with more precise fixations than guesses in all time bins except for 0.5–1s (see Figure 4a).

**Figure 4.**
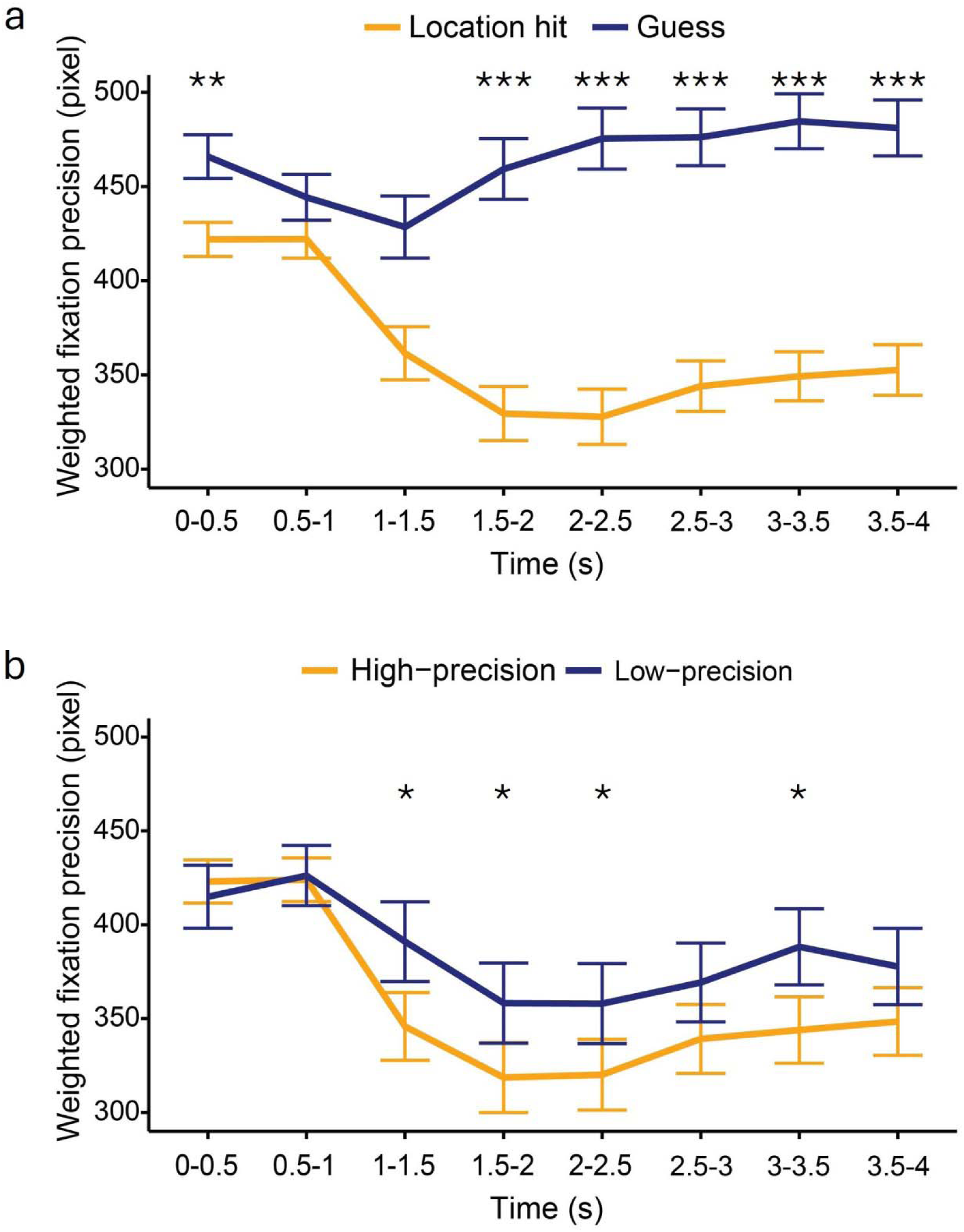
Duration-weighted fixation precision for (a) location hits vs. guess trials and (b) high vs. low-precision trials. Error bars depict standard errors of the means derived from the linear mixed-effects models. * p< 0.05; ** p < 0.01; *** p< 0.001. All significant values were FDR-corrected across all time bins.

**Table 2.** Results from linear mixed effects models employing trial type and time bin as predictors of fixation precision.

|  | df | F | p |
| --- | --- | --- | --- |
| *Location hit vs. guess* |  |  |  |
| Trial type | (1, 8658) | 390.07 | < 0.001 |
| Time bin | (7, 8643) | 3.24 | 0.002 |
| Trial type x time bin | (7, 8638) | 10.75 | < 0.001 |
| *High vs. low-precision* |  |  |  |
| Trial type | (1, 5194) | 9.46 | 0.002 |
| Time bin | (7, 5186) | 7.13 | < 0.001 |
| Trial type x time bin | (7, 5184) | 1.47 | 0.172 |

As is also shown in Table 2, the model contrasting high vs. low precision trials yielded significant main effects of trial type and time bin, indicative of more precise fixations for high than for low-precision trials, and for later than for earlier time bins (Figure 4b). Analyses following up the main effect of time bin revealed lower fixation precision in the first two bins than the later bins starting from the bin of 1.5-2s (ps < 0.05, FDR-corrected). Despite the non-significant trial type x time bin interaction, we conducted an exploratory analysis for each time bin to compare fixation precision according to trial type. Significant differences were identified from 1–1.5s post-cue until 2–2.5s. A significant difference between the two trial types was also evident for the 3-3.5s bin (Figure 4b).

The analyses conducted on the pre-stimulus time bin (-0.5 – 0s) did not reveal a significant difference between location hits and guesses (p = 0.799), or between high vs low precision trials (p = 0.268).

### Fixation pattern similarity analyses

#### Effects of retrieval success and memory precision on pattern similarity

Figure 5a displays the fixation density maps associated with location hits and guesses, as well high and low precision memory judgments, aggregated across all trials and participants. To facilitate comparison, the studied location associated with all trials across participants were realigned to the same location, and their corresponding fixation locations were adjusted accordingly. As is evident from the figure, fixations were more densely distributed around the studied location for location hits than for guesses, and for high-precision than low-precision trials.

**Figure 5.**
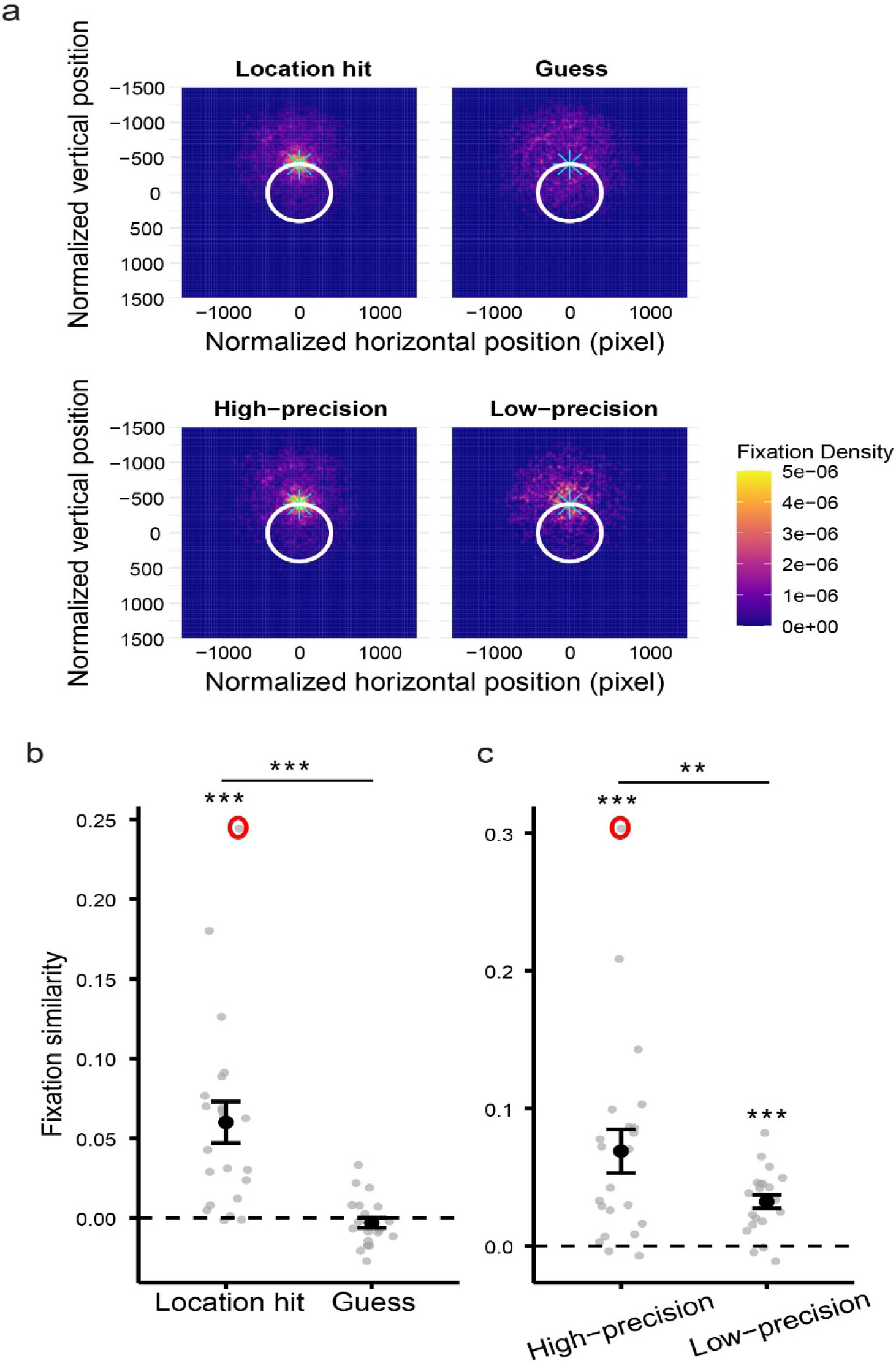
a: Heatmaps illustrating the density of eye fixations (weighted by fixation duration) around the studied location (marked with the cyan star) for different trial types. To facilitate comparison, all studied locations were realigned to the top point of the circle, and the corresponding fixation locations were adjusted accordingly. b: Estimates of fixation similarity of location hits and guesses. c: Estimates of fixation similarity of high and low-precision trials. *** p < 0.001. ** p < 0.01. Error bars represent the standard errors of the mean. Excluding the data from the participant with the outlying data point (circled, identified as >3 standard deviations from the mean) did not alter the results.

A pairwise t-test comparing the similarity metrics (see the section of ‘Pattern similarity analysis’) between trial types revealed greater across-trial similarity for location hits than for guesses [t(21) = 4.66, p < 0.001, Cohen’s d = 1.43], and for high-precision than for low-precision judgments [t(21) = 2.89, p = 0.009, Cohen’s d = 0.45]. The similarity metrics for all trial types other than guesses were significantly greater than 0 (for guesses, p = 0.348; for other trial types, ps < 0.001, see Figures 5b and c).

#### Fixation pattern similarity predicts precision of memory judgments

We went on to examine whether across-trial fixation similarity was predictive of the precision of location hits. As is shown in Table 3, the relationship between fixation similarity and memory precision was significant, with greater similarity associated with lower distance error and thus higher memory precision (see Figure 6a). In a complementary analysis, participant-level pairwise correlations were computed between the trial-wise metrics of absolute distance error and fixation pattern similarity, and the mean Fisher z transformed correlation coefficients are shown in Figure 6b. A one-sample t-test revealed that the mean correlation coefficient was significantly below 0 [t(21) = 3.87, p < 0.001, Cohen’s d = 0.83], consistent with the findings from the LME analysis.

**Figure 6.**
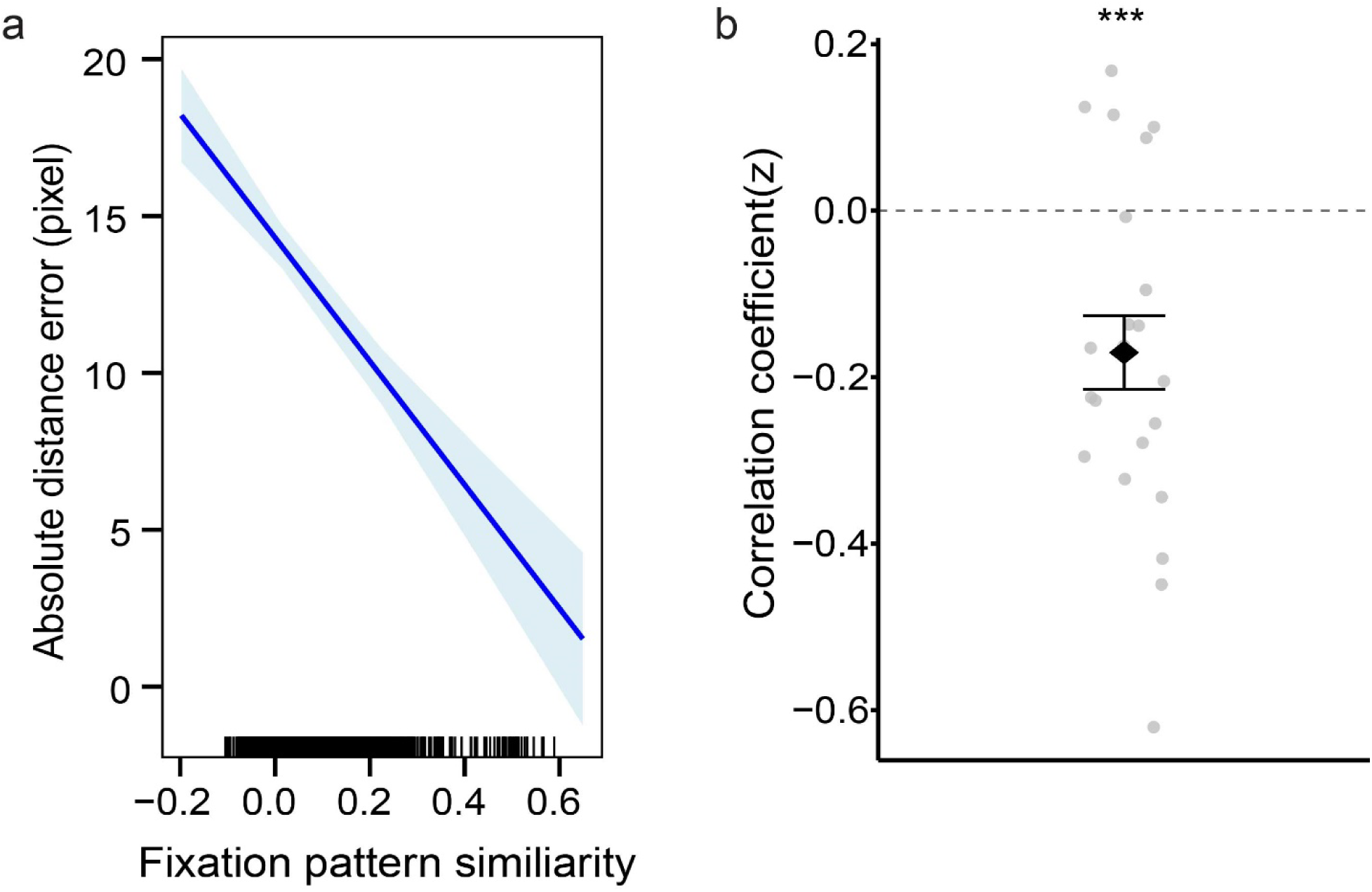
a: Plot depicting the relationship between trial-wise fixation pattern similarity and the model fitted values of judgment precision for location hits, as derived from the LME model employing fixation pattern similarity as the predictor of judgment precision. The shaded areas reflect the 95% confidence intervals. b: Participant-wise Pearson correlation coefficients (Fisher z-transformed) for the relationship across trials between fixation pattern similarity and judgment precision. The error bar represents the standard error of the mean. *** p < 0.001.

**Table 3.**
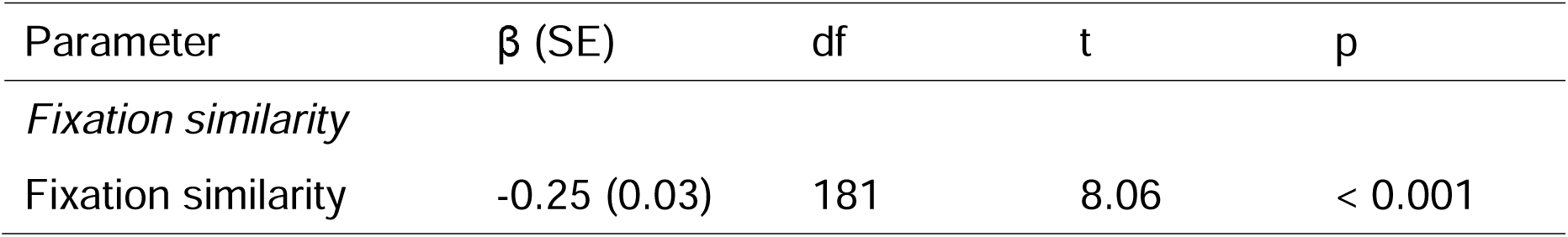
Results from linear mixed models employing fixation similarity as a predictor of trial-wise memory precision. Standardized regression coefficients (β) are reported.

| Parameter | $\beta$ (SE) | df | t | p |
| --- | --- | --- | --- | --- |
| *Fixation similarity* |  |  |  |  |
| Fixation similarity | -0.25 (0.03) | 181 | 8.06 | < 0.001 |

#### Control analysis: fixations in reference to the judged location

To determine whether the findings reported above merely reflected preparation for the upcoming location judgment rather than reflecting retrieved memory content, we examined the mean Euclidean distance of all fixations relative to the judged location as a function of trial type. The fixation to judged location distance was significantly closer for location hits (M = 354, sd = 71) than guess trials [M = 399, sd = 65), t(21) = 3.44, p = 0.002, Cohen’s d = 0.73]. There was, however, no significant difference between high (M =357, sd = 82) and low-precision trials (M= 352, sd = 68), p = 0.679.

We also analyzed the similarity of the fixation patterns associated with different trial types relative to the judged location (see Supplemental Figure 1 for the heatmaps illustrating the density of eye fixations around the judged location for different trial types). Pairwise t-tests revealed that across-trial similarity was higher for location hits than for guesses [t(21) = 2.82, p = 0.010, Cohen’s d = 1.08]. By contrast, similarity did not differ between high and low-precision judgments (p = 0.338). The similarity metrics for all trial types other than guesses were significantly greater than 0 (for guesses, p = 0.105; for other trial types, ps < 0.008).

#### Hippocampal BOLD signals predict precision of memory judgments

In light of prior findings that fMRI BOLD activity in the hippocampus and the left AG independently predicted trial-wise memory precision (Hou et al., 2025b), we examined the relationships between fixation precision, precision of location judgments and neural activity in these regions. As shown in Table 4, trial-wise BOLD activity in the hippocampus (see Figure 7a) was predictive of the precision of location judgments. We went on to conduct a pairwise correlation analysis between hippocampal BOLD activity and judgement precision across participants. As is illustrated in Figure 7b, the mean pairwise correlation coefficient across participants was significantly below 0 [t(21) = 2.36, p = 0.028, Cohen’s d = 0.50], consistent with the findings from the LME analysis. By contrast, the association between trial-wise AG activity and judgment precision only approached significance. We therefore focused on hippocampal activity in the subsequent analyses.

**Figure 7.**
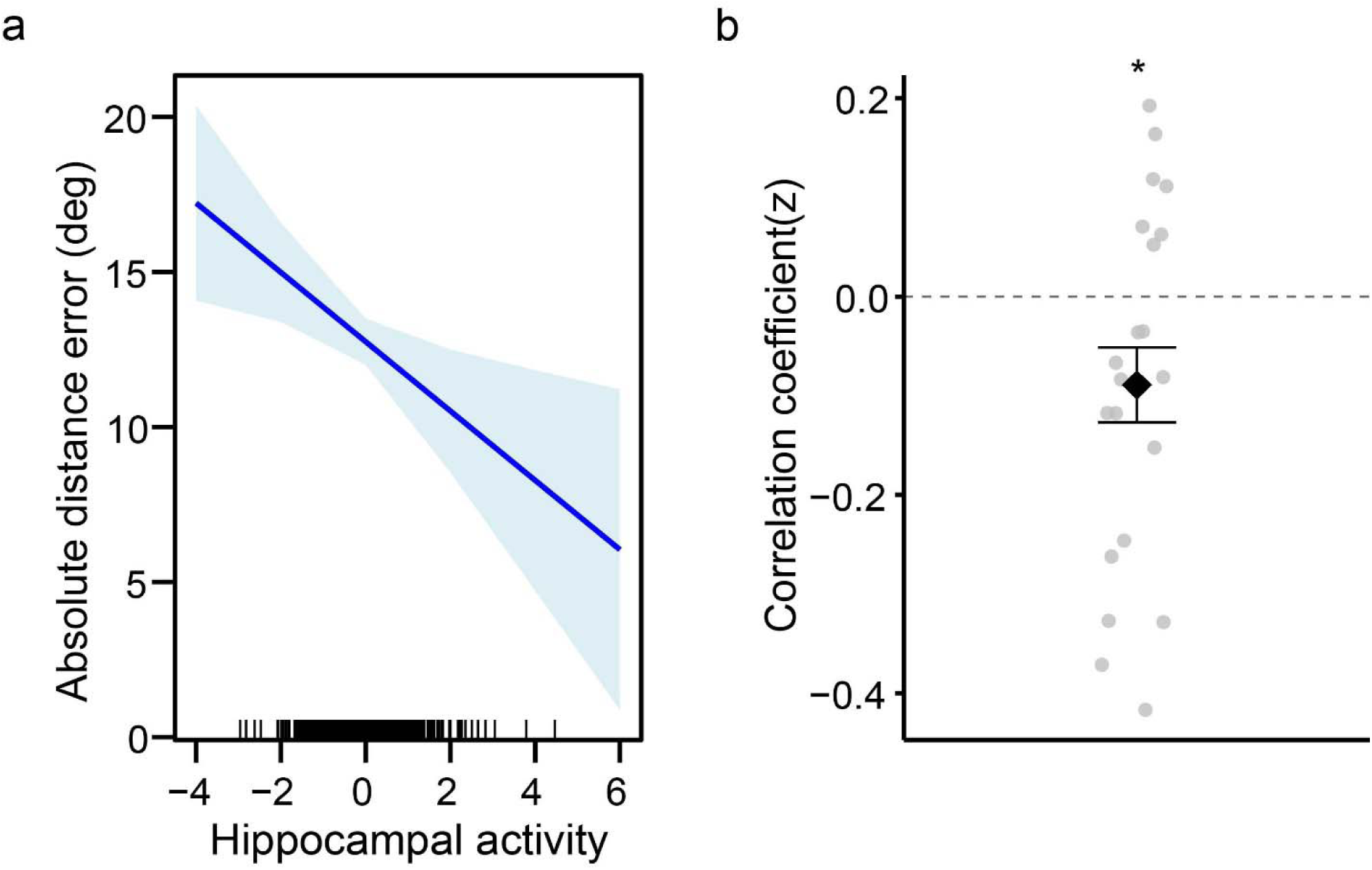
a: Plot depicting the relationship between trial-wise hippocampal BOLD activity and the model fitted values of memory precision for location hits, as derived from the LME model employing hippocampal activity as the predictor of precision. The shaded areas reflect the 95% confidence intervals. b: Participant-wise Pearson correlation coefficients (Fisher z-transformed) for the relationship across trials between hippocampal activity and judgment precision. The error bar represents the standard error of the mean. * p < 0.05.

**Table 4.**
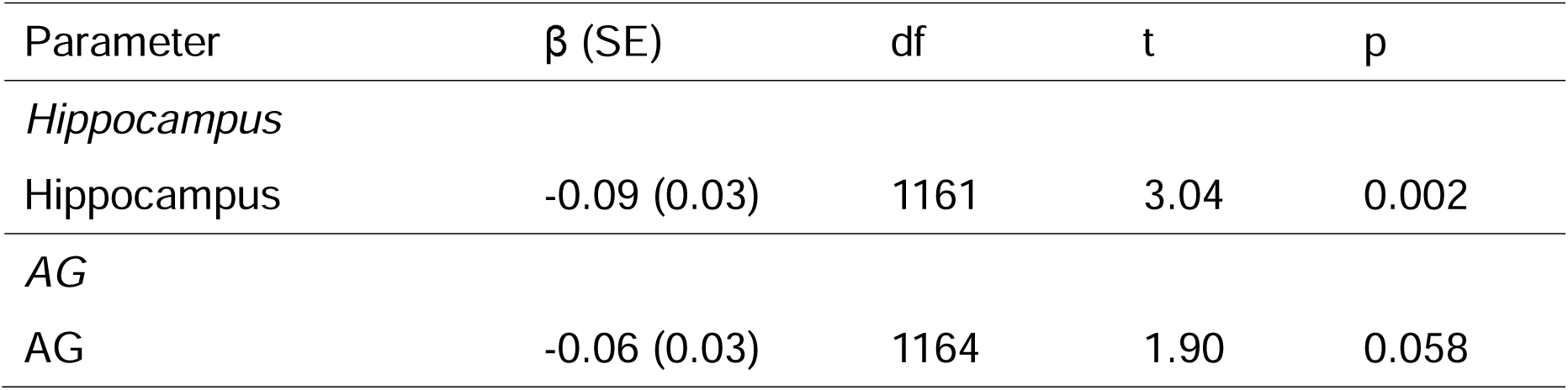
Results from linear mixed models examining the relationships between BOLD activity in the hippocampal and AG ROIs and memory precision. Standardized regression coefficients (β) are reported.

#### Relationships among memory precision, fixation metrics and hippocampal activity

Lastly, we assessed whether eye fixation metrics and hippocampal BOLD activity independently predicted judgment precision. As is evident in Table 5, when both fixation precision and fixation pattern similarity were included in the same model as predictors of the precision of location judgments, only similarity remained significant. As is also shown in the table, each fixation metric independently predicted judgment precision when hippocampal activity was included in the model. Moreover, both fixation pattern similarity and hippocampal activity remained significant predictors after controlling for fixation precision.

**Table 5.**
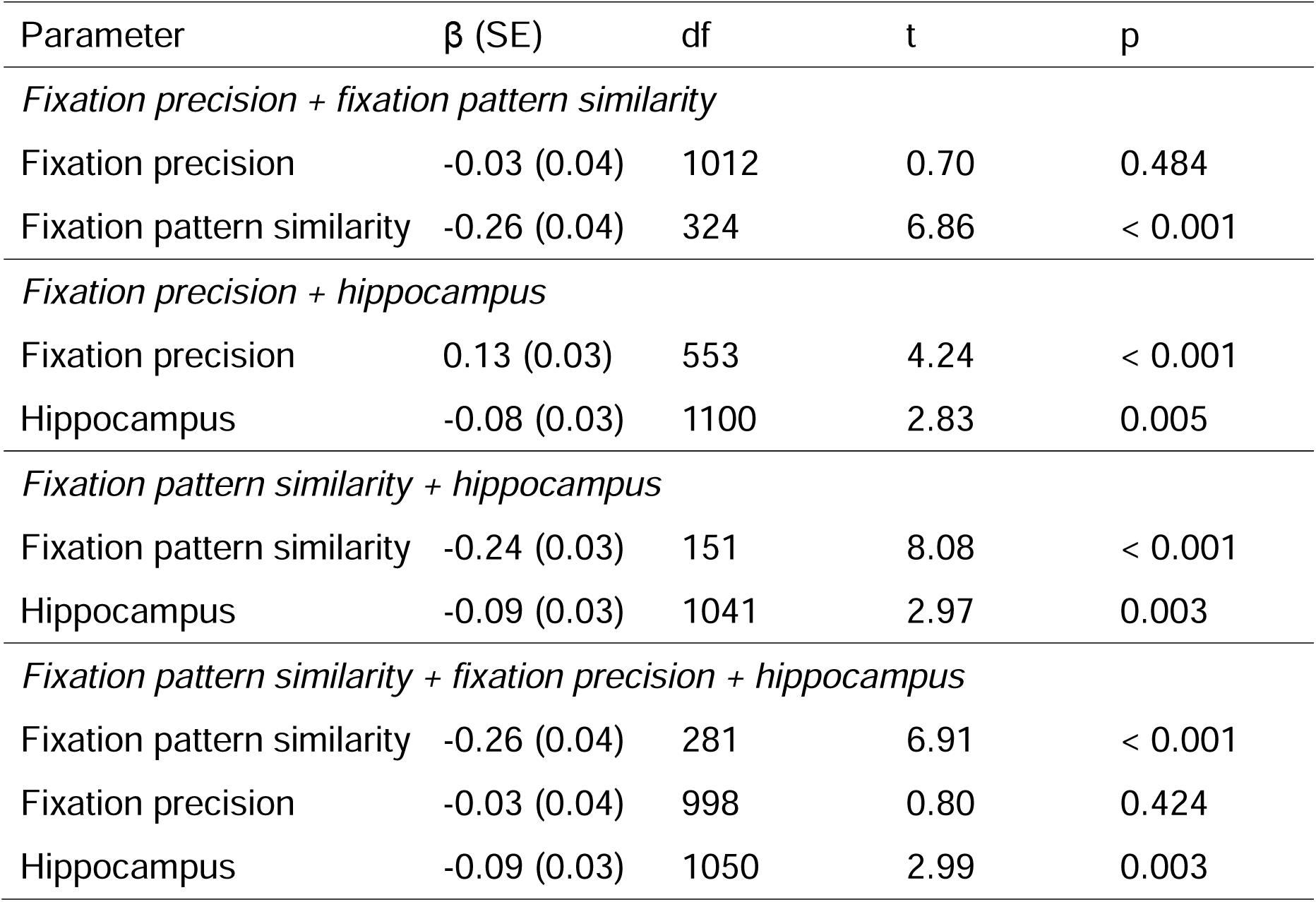
Results from linear mixed models employing fixation precision, fixation pattern similarity and neural activity in the hippocampus as predictors of trial-wise memory precision. Standardized regression coefficients (β) are reported.

| Parameter | $\beta$ (SE) | df | t | p |
| --- | --- | --- | --- | --- |
| *Fixation precision + fixation pattern similarity* |  |  |  |  |
| Fixation precision | -0.03 (0.04) | 1012 | 0.70 | 0.484 |
| Fixation pattern similarity | -0.26 (0.04) | 324 | 6.86 | < 0.001 |
| *Fixation precision + hippocampus* |  |  |  |  |
| Fixation precision | 0.13 (0.03) | 553 | 4.24 | < 0.001 |
| Hippocampus | -0.08 (0.03) | 1100 | 2.83 | 0.005 |
| *Fixation pattern similarity + hippocampus* |  |  |  |  |
| Fixation pattern similarity | -0.24 (0.03) | 151 | 8.08 | < 0.001 |
| Hippocampus | -0.09 (0.03) | 1041 | 2.97 | 0.003 |
| *Fixation pattern similarity + fixation precision + hippocampus* |  |  |  |  |
| Fixation pattern similarity | -0.26 (0.04) | 281 | 6.91 | < 0.001 |
| Fixation precision | -0.03 (0.04) | 998 | 0.80 | 0.424 |
| Hippocampus | -0.09 (0.03) | 1050 | 2.99 | 0.003 |

Additional LME analyses revealed that fixation pattern similarity was significantly associated with fixation precision (β = -0.61, SE = 0.03, t = 22.39, p < 0.001). However, hippocampal activity did not predict either fixation metric (ps > 0.104).

## Discussion

The present study examined the relationship during retrieval between eye fixations and memory precision for the spatial location of studied object images. We identified robust associations between the ‘precision’ of fixations occurring prior to an overt memory judgment and both the success and precision of location memories. The associations took the form of fixations that were closer to the studied location for memory judgments reflecting successful location retrieval than for guesses (a retrieval success effect), and for judgments associated with high vs. low precision (a memory precision effect). Additionally, across-trial similarity between fixation patterns was higher for memory judgments associated with successful location retrieval than for guess trials, and for high- than for low-precision trials. Time series analyses revealed that the effect of retrieval success on fixation precision persisted for almost the entire recall phase. The effects of memory precision emerged around 1-1.5s after the onset of the cue. Lastly, fixation precision and pattern similarity each predicted trial-wise memory precision independently of hippocampal fMRI BOLD activity. Below, we discuss these findings in more detail.

Consistent with prior studies (Buchel et al., 2024; Hannula et al., 2009; Johansson & Johansson, 2014; Schmidig et al., 2025), we observed higher fixation precision prior to overt memory judgments based on retrieved location information compared to judgments associated with guesses. More importantly, fixation precision tracked the precision of overt location judgments on a trial-wise basis. That is, eye fixations were closer to the studied location prior to the retrieval of precise rather than less precise location memories. These findings extend prior work (e.g., Buchel et al., 2024; Schmidig et al., 2025) by indicating that eye fixations are predictive not only of the likelihood of successful retrieval, but also of the fidelity of the retrieved mnemonic content.

The retrieval success effect on fixation precision persisted across nearly the entire recall phase. By contrast, a precision effect emerged around 1-1.5s after the image onset. These results suggest that the sensitivity of fixations to memory content is present relatively early during recall and remains stable thereafter. The reason for the latency difference between the two effects is unclear. One possibility is that the experimental design did not provide sufficient support for participants to accurately fixate on the studied location during the early stages of recall. As is evident from Figure 1, the test image was presented in the absence of the reference circle for the first second of the recall phase. Consequently, participants would have been unable to use the circle as a guide for fixations until some time after the onset of the retrieval cue. Alternatively, the longer latency of the precision effect might indicate that the processes supporting memory precision operate more slowly than those supporting retrieval success. It should be noted, however, that the low temporal resolution of these analyses (500-ms time bins), combined with the limited number of high and low precision trials contributing to each bin, prevented a detailed characterization of the time-course of these effects. Some information about the timing of the retrieval processes associated with differing levels of memory precision is available however from the findings of two prior event-related potential (ERP) studies (Murray et al., 2015; Ladyka-Wojcik et al., 2025). In both cases, memory precision was associated with modulation of the amplitude of the so-called ‘left parietal effect’, a retrieval-related ERP modulation that is typically observed between about 500-800ms after cue onset (for a recent review, see Kwon et al., 2023). It is unclear how the present findings regarding the precision of eye fixations might relate to these ERP findings. Future studies with greater sensitivity and temporal resolution are needed to better characterize the temporal dynamics of retrieval-related fixations and to examine their relationship with corresponding electrophysiological effects.

While fixation patterns were randomly distributed across guess trials, they were significantly correlated across trials on which participants successfully retrieved location information, and these correlations increased with memory precision. These findings suggest that retrieval of high-fidelity location information is associated with a systematic and relatively stereotypical spatial distribution of fixations. Of interest, despite the strong across-participant correlation between fixation pattern similarity and fixation precision (r = 0.88), when both metrics were employed in the same model as predictors of the precision of overt memory judgments, only pattern similarity remained significant. While it remains to be established whether these findings generalize to memory features other than location, our results highlight the importance of examining the spatial distribution of fixations when assessing the relationship between eye movements and memory retrieval, rather than relying solely on a univariate metric such as mean Euclidian distance.

The present findings are consistent with the proposal that eye fixations might serve as a covert or indirect index of memory retrieval (Schmidig et al., 2025). However, to employ fixations as a marker of retrieval, it is necessary to demonstrate that they are selectively associated with memory judgments that are reflective of successful memory retrieval. To address this question, we compared fixation precision and pattern similarity relative to the judged rather than the studied location. Fixations were closer to the judged location for location hits than for guess trials, and fixation patterns associated with location hit trials were consistently clustered around the judged location, whereas guesses were associated with more diffusely distributed patterns. Thus, gaze patterns did not track judged location when there was an apparent failure to retrieve location information (this null finding is especially notable given that the use of a sample-wide cut-off raises the possibility that in some participants a proportion of the trials were mis-categorized as guesses rather than location hits). That is, the findings indicate that eye fixations were selectively predictive of judged location only when retrieval was successful.

We also examined the relationships among precision-related fixations, neural activity and the precision of memory judgments. This analysis was motivated by our prior findings from a dataset that included the current sample as a sub-sample (Hou et al., 2025b), where it was reported that fMRI BOLD signals in the hippocampus and left AG were each predictive of memory precision. In the reduced data set available here only hippocampal activity tracked memory precision, consistent the proposal that this structure contributes to the encoding and reinstatement of high-precision mnemonic information (Ekstrom & Yonelinas, 2020). By contrast, we were unable to identify a relationship between AG activity and memory precision. This null finding likely reflects reduced power due to the smaller sample size of the present study than that of the parent study. It is worth noting however that in a subregion of the AG ROI employed by Hou et al. (2025a) we were able to identify a significant relationship between AG activity and judgment precision (Richter ROI, β = -0.08, SE = 0.03, t = 2.72, p = 0.007).

Although both fixation precision and pattern similarity were predictive of the precision of the subsequent location judgment, neither variable was significantly correlated with hippocampal activity. Moreover, when included in the same model, the fixation metrics and hippocampal activity independently predicted judgment precision. Among other possible explanations, this finding might indicate that some neural variable other than univariate fMRI BOLD activity reflects the contribution of the hippocampus to our fixation precision and pattern similarity metrics (e.g., a quite different finding might have emerged had we employed electrophysiological methods to sample hippocampal activity; cf. Kragel & Voss, 2022). Alternately, of course, it might be that the fixation precision and pattern similarity metrics reflect a non-hippocampal mechanism.

## Limitations

The present study has several limitations. First, we only assessed location memory, leaving it unresolved whether the relationships between eye fixations and memory precision identified here extend to other contextual features. Second, the experimental design was not optimized to characterize the time course of the relationship between fixations and memory precision. Third, the low spatial resolution of the fMRI data precluded the opportunity to investigate relationships between precision-related eye movements and fMRI BOLD activity at the level of hippocampal subfields. These limitations can all be addressed in future research.

## Conclusion

The present findings indicate that eye fixations are closer to and more tightly clustered around a studied location when a location memory was retrieved with high rather than low precision. Thus, the findings are consistent with the proposal that eye fixations track the fidelity of retrieved mnemonic content.

## Data availability statement

The data that support the findings of this study are available from the authors on request subsequent to a formal data sharing agreement. The software, functions and syntax of the models used to generate the results are specified in the Methods & Materials section.

## Author contribution

Mingzhu Hou: Conceptualization; Data curation; Formal analysis; Investigation; Methodology; Software; Validation; Visualization; Writing—Original draft; Writing—Review & editing. Luke R. Pezanko: Investigation; Writing—Review & editing. Sabina Srokova: Methodology; Writing—Review & editing. Paul F. Hill: Writing—Review & editing. Arne D. Ekstrom: Funding acquisition; Writing—Review & editing. Michael D. Rugg: Conceptualization; Funding acquisition; Methodology; Resources; Supervision; Writing—Original draft; Writing—Review & editing.

## Acknowledgments

This work was supported by the National Institute of Neurological Disorders and Stroke (grant number R01NS114913). We thank our experimental participants for volunteering their time.

