## supplemental materials for "Retrieval-related eye movements are predictive of memory precision"

Supplemental Figure 1 shows the heatmaps reflecting the density of fixations around the judged location associated with different trial types. As is evident from this figure, fixations were more tightly clustered around the response location for location hits than for guess trials. By contrast, high and low-precision trials showed similar fixation patterns.

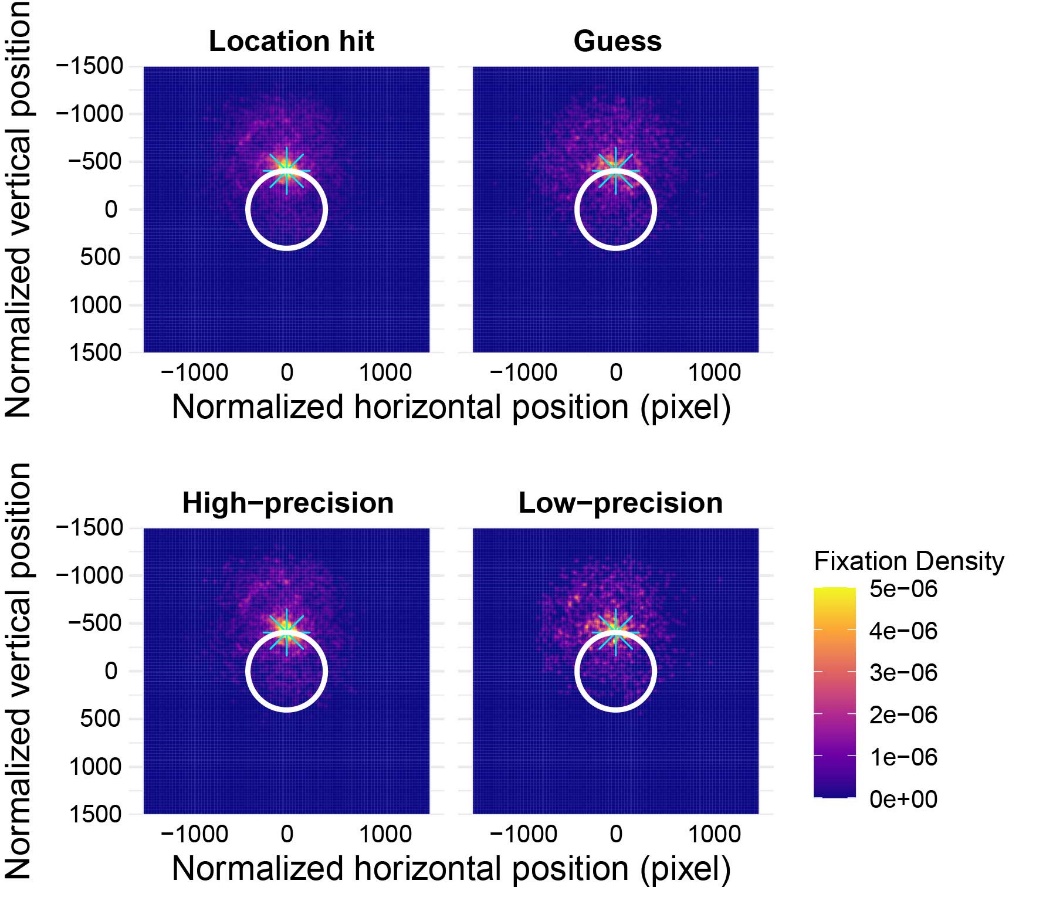

Supplemental Figure 1. Heatmaps illustrating the density of eye fixations (weighted by fixation duration) around the judged location (marked as the cyan star) for different trial types. To facilitate comparison, all response locations were realigned to the top point of the circle, and the corresponding fixation locations were adjusted accordingly.
